# Development of a tryptophan-based dual selection system reveals the spatial organization of S-layer assembly during cytokinesis in *Sulfolobus acidocaldarius*

**DOI:** 10.64898/2026.06.29.735232

**Authors:** Sherman Foo, Buzz Baum

## Abstract

*Sulfolobus acidocaldarius* is a thermoacidophilic archaeon used as a model system for studying fundamental cellular processes and for emerging biotechnological applications. However, the limited availability of selectable markers restricts advanced genetic manipulation in this organism. Here, we report the development of a tryptophan auxotrophy-based selection system in *S. acidocaldarius*. A Δ*trpBA* mutant was constructed in the Δ*pyrE* background strain using a classical pop-in/pop-out recombination strategy. The resulting mutant exhibited little growth defects in rich medium, likely due to exogenous tryptophan supplied by complex nutrients, but failed to grow in a newly developed defined Brock-based amino acid dropout medium lacking tryptophan. Exploiting both uracil and tryptophan auxotrophies, we achieved dual-plasmid co-transformation and co-expression of the surface layer proteins and a dominant-negative mutant of the AAA-ATPase Vps4, revealing that the accumulation of surface layer lattice forming protein SlaA at the midzone of division-arrested cells together with its membrane anchor SlaB. Together, these results provide evidence for spatial regulation of S-layer assembly during archaeal cytokinesis while expanding the genetic toolkit available for *S. acidocaldarius*.

**Importance:** *Sulfolobus acidocaldarius* is a key archaeal model organism for studying cellular processes shared with more complex life and is increasingly used for biotechnological applications. Here, we establish tryptophan auxotrophy as a new selectable marker in *S. acidocaldarius*, expanding the range of genetic selection systems available in this organism. By developing a defined Brock-based dropout medium, we enable stringent amino acid auxotrophy selection and precise control over nutrient composition. This system can be combined with existing uracil-based selection to support dual auxotrophy workflows, enabling co-transformation, simultaneous expression of multiple proteins, and more sophisticated genetic manipulation strategies. Using both markers, we show that S-layer proteins are localised to the division bridge in cytokinesis-arrested cells. This exemplifies ways in which the expanding molecular genetic tool kit available for *Sulfolobus acidocaldarius* is furthering our understanding of archaeal cell biology.

## Introduction

Since their discovery in 1972 (1), members of the *Sulfolobales* have become prominent archaeal model systems for studying fundamental biological processes conserved across evolution (2). As currently the closest genetically tractable archaeal relatives of eukaryotes, the *Sulfolobales* have provided key insights into processes such as cell division and DNA segregation (3), and have illuminated our understanding of eukaryogenesis (4).

Despite these advances, genetic manipulation in *Sulfolobus acidocaldarius* remains constrained by the limited number of available selectable markers. Current systems rely primarily on uracil auxotrophy (e.g. *pyrEF* or *pyrB* selection) (5, 6), limiting more complex genetic studies requiring multiplexed gene expression. More recently, agmatine auxotrophy based on the deletion of the arginine decarboxylase have been described in both *S. acidocaldarius* and the related *Saccharolobus islandicus* (7, 8), alongside antibiotic-based selection systems such as simvastatin resistance in *Sc. islandicus* (9). However, no antibiotic-based selection system has been described for *S. acidocaldarius* thus far, limiting the options available in this organism. Compared to the closely related *Sc. islandicus*, *S. acidocaldarius* is faster growing and possesses a comparatively stable genome with fewer active mobile genetic elements (10), contributing to its robustness as a genetically tractable model archaeon. Expanding the repertoire of selectable markers in this organism is therefore especially important for enabling increasingly sophisticated genetic and cell biological studies.

Amino acid auxotrophy-based selection systems offer a robust and versatile alternative to antibiotic-based approaches, and are widely used in model organisms such as *Escherichia coli* and *Schizosaccharomyces pombe*. Recent efforts to develop defined cultivation media for *S. acidocaldarius* have focused on improving growth characteristics and reducing costs in order to facilitate biotechnological applications through increased control over nutrient composition (11); however, these formulations have not been widely adapted for the selective growth of auxotrophic mutants thus far. Here, we address this limitation by developing a defined Brock-based dropout medium that enables stringent selection of amino acid auxotrophies. Using this medium, we establish tryptophan auxotrophy as a robust selectable marker through deletion of the *trpBA* tryptophan synthase genes.

Furthermore, we demonstrate the utility of this system by constructing a complementation vector and by combining tryptophan and uracil auxotrophy for dual selection. This approach enables the co-transformation and maintenance of two independently selectable plasmids, facilitating multiplexed gene expression, combinatorial protein tagging, and more flexible experimental workflows. Using this platform, we investigate the localization of S-layer proteins during cytokinesis and identify the co-enrichment of SlaA and SlaB at the division midzone of division-arrested cells. Together, these results expand the genetic toolkit available for *S. acidocaldarius* and exemplify the way these tools can be used to further our understanding of the molecular mechanisms underlying archaeal cell organization and cell division.

## Results

### Generation of a tryptophan auxotrophic mutant of *S. acidocaldarius*

Tryptophan biosynthesis is a multistep process utilizing chorismate, the end product of the shikimate pathway (Fig. S1). In *S. acidocaldarius*, the genes involved in tryptophan biosynthesis are organized in an operon (*saci1421*- *saci1427*) (10), with the final catalytic steps carried out by tryptophan synthase, composed of the TrpB and TrpA subunits (Fig. S1).

To construct a tryptophan auxotrophic *S. acidocaldarius*, we targeted deletion of the tryptophan synthase genes *trpBA* (*saci1421* and *saci1422*), thereby blocking the final step of tryptophan biosynthesis while avoiding disruption of upstream intermediates. Gene deletion was performed using the classical pop-in/ pop-out method based on uracil auxotrophy and 5-fluoroorotic acid (5-FOA) counterselection in the Δ*pyrE* MW001 background strain (12). Successful deletion of the *trpBA* genes was confirmed by PCR genotyping (Fig. 1A).

**Figure 1.**
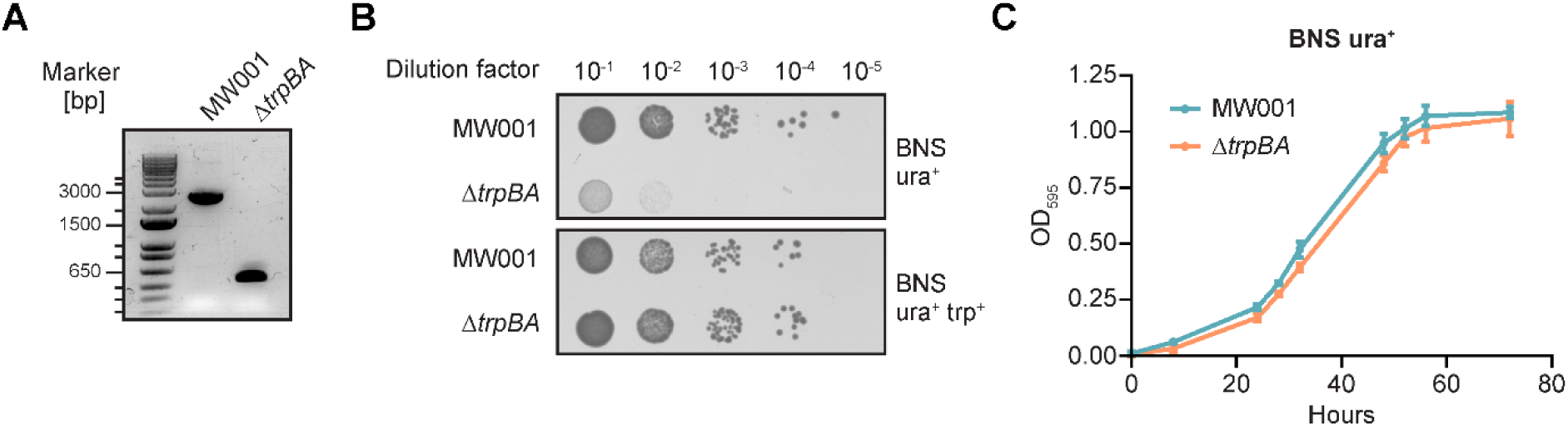
The tryptophan synthase Δ*trpBA* mutant of *S. acidocaldarius*. **(A)** PCR genotyping utilizing flanking primers confirms the deletion of the 2009bp Δ*trpBA* genes (*saci1421* and *saci1422*) in the Δ*pyrE* MW001 background strain. **(B)** Spot assay of MW001 and Δ*trpBA* on rich BNS medium shows high background growth of the Δ*trpBA* mutant in the absence of tryptophan supplementation. **(C)** Growth of the Δ*trpBA* mutant in rich liquid BNS medium is indistinguishable from MW001 even in the absence of tryptophan supplementation. Error bars represent mean ±SD.

As evident by our successful isolation of the Δ*trpBA* mutant on standard **B**rock medium supplemented with **N**Z-amine, **S**ucrose and uracil (hereafter referred to as BNS), tryptophan supplementation was not required to support growth under these conditions. This was confirmed by a spot assay, where high background growth of the Δ*trpBA* mutant can be observed in the absence of tryptophan supplementation (Fig. 1B). Furthermore, growth analysis in liquid BNS further confirms little significant difference between the growth of the Δ*trpBA* mutant and the MW001 background strain in the absence of tryptophan supplementation (Fig. 1C). Taken together, our results indicate that optimization of the rich BNS culture medium is required for the use of tryptophan auxotrophy as a robust selection marker in *S. acidocaldarius*.

### Development of a defined dropout medium for isolation of auxotrophic mutants

While a defined culture medium for *S. acidocaldarius* have been previously reported (11), Brock medium remains the gold standard for laboratories worldwide (13–17). This basal medium is typically supplemented with a protein-based complex substrate such as NZ-amine, yeast extract or tryptone as a source of essential amino acids, peptides and other nutrients. Additionally, a secondary carbon source, typically sucrose or dextrin, is included to support growth. As NZ-amine represents the only complex and undefined component of our BNS medium, we sought to replace it with a defined amino acid mixture while maintaining all other components unchanged, thereby minimizing alterations to media composition and preparation.

Most commercially available NZ-amine and tryptone formulations lack detailed information regarding their amino acid composition, consistent with their undefined nature. However, NZ-Amine A (Sigma-Aldrich, C0626) provides an approximate amino acid profile, which we used as a basis for formulating a defined amino acid mixture.

This, together with prior analyses of related complex substrates (11), enables the formulation of a defined amino acid mix (Fig. 2A) to replace the complex component in BNS medium. We term this Brock dropout medium (hereafter referred to as BDM). Spot assay and growth analysis revealed that the growth of the Δ*trpBA* mutant was completely abolished on BDM (Fig. 2B, C), and was only possible upon the addition of at least 10μg/mL tryptophan (Fig. 2D).

**Figure 2.**
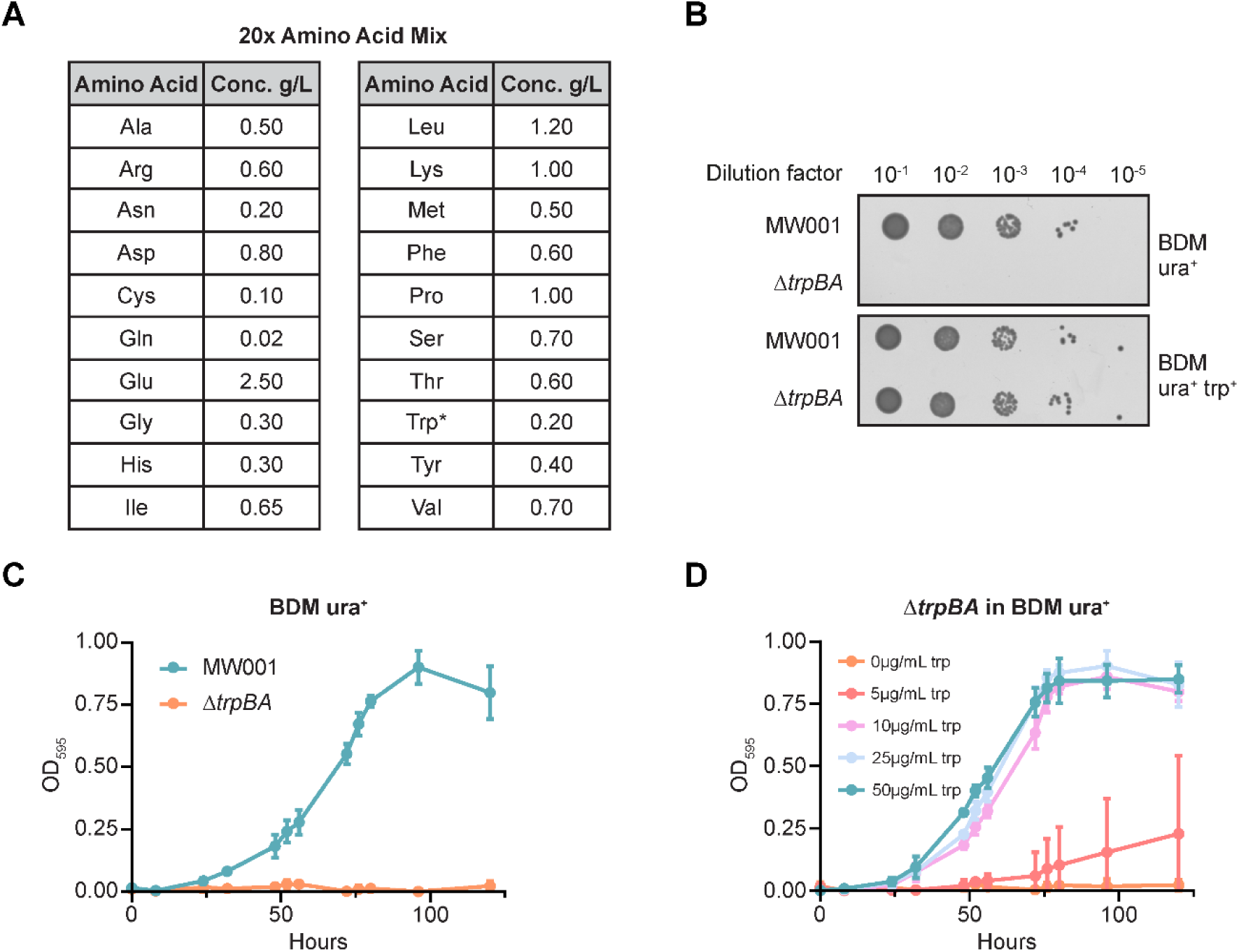
Stringent selection of tryptophan auxotrophy in defined dropout medium. **(A)** Amino acid mix (20x stock) formulated based on analysis of amino acid composition of NZ-amine A (Sigma-Aldrich, C0626) and other related substrates (11). *Tryptophan is omitted from the mix in this study. **(B)** Spot assay showing tight selection of the Δ*trpBA* mutant on BDM, with no background growth in the absence of tryptophan supplementation. Growth curves of **(C)** MW001 and the Δ*trpBA* mutant in BDM, and **(D)** the Δ*trpBA* mutant with tryptophan supplementation in BDM. Error bars represent mean ±SD.

To further characterize growth in BDM, the MW001 parental strain was grown and analysed under these conditions. MW001 cells grown in BDM exhibited similar profiles of the ESCRT-III homologs CdvB, CdvB1, and CdvB2 compared to those grown in BNS, as determined by flow cytometry (Fig. S2A). Distinct populations corresponding to low (1N) and high (2N) DNA content could be resolved, and cell cycle-dependent changes in ESCRT-III protein levels in asynchronous populations were consistent with previous reports (18, 19).

Growth in BDM was however characterized by a longer lag phase in contrast to BNS (Fig. 1C, 2C), and was only modestly improved in the linear phase by the addition of Wolin’s vitamin solution (20) (Fig. S2B). To assess protein expression under defined conditions, we expressed the surface layer (S-layer) protein SlaA-HA (21) under an arabinose-inducible promoter using the pSVAaraFX-HA plasmid (*pyrEF* selection) (22). Induction with arabinose for 4 hours resulted in successful expression of SlaA-HA in both BNS and BDM (Fig. S2C), with localization to the cell surface as expected in both conditions (Fig. S2D). However, overall protein expression levels were markedly higher in rich BNS medium compared to defined BDM both before and after induction (Fig. S2C).

Having successfully developed a stringent defined selection medium for amino acid auxotrophy in *S. acidocaldarius*, we next generated a Δ*trpF* anthranilate isomerase mutant (Fig. S3A, B) to test if the counterselection agent 5-fluoroanthranilic acid (5-FAA) could also be utilized in this organism. When spotted on plates containing the toxic antimetabolite 5-FAA (23), none of the strains tested were able to survive (Fig. S3C), suggesting that deletion of *trpF* was insufficient to prevent the formation of toxic fluorinated tryptophan derivatives from 5-FAA under these conditions. Therefore, unlike in other model organisms (23), 5-FAA does not appear to function as an effective counterselection agent for the tryptophan biosynthetic pathway in *S. acidocaldarius* under the conditions tested here.

### Complementation of tryptophan auxotrophy in *S. acidocaldarius*

We next turned our attention to the design of a complementation vector to rescue tryptophan auxotrophy. Using the pSVAaraFX-HA plasmid backbone, we replaced the *pyrEF* cassette with the *trpBAD* genes from *Saccharolobus solfataricus* (*sso0888* - *sso0890*). The resulting construct, termed pSF251 (Fig. 3A), was transformed into the Δ*trpBA* mutant.

**Figure 3.**
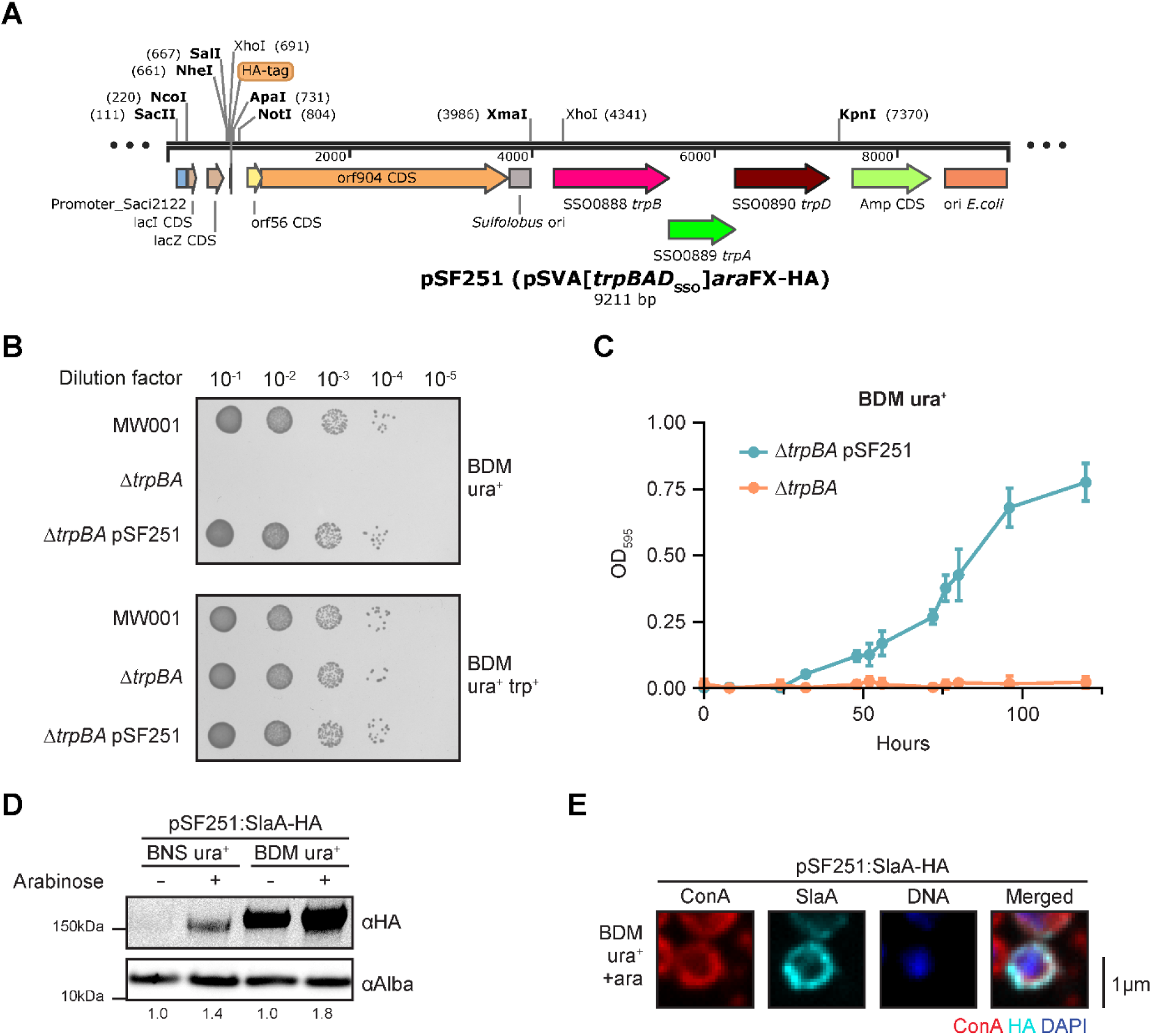
Complementation of tryptophan auxotrophy in *S. acidocaldarius*. **(A)** Plasmid map of the tryptophan complementation vector pSF251, based on the pSVAara-FX-HA plasmid (22). **(B)** Spot assay showing complementation of the Δ*trpBA* mutant with pSF251, restoring its growth on defined BDM without tryptophan supplementation. **(C)** Growth curves of the Δ*trpBA* mutant with or without pSF251 complementation vector, in BDM without tryptophan supplementation. Error bars represent mean ±SD. **(D)** Western blot of SlaA-HA expression from pSF251 with or without 4h of arabinose induction. The DNA-binding protein Alba was used as a loading control. Values indicate the normalised HA signal intensity expressed relative to the corresponding uninduced sample. **(E)** Representative single-plane confocal micrographs of Δ*trpBA* in BDM expressing SlaA-HA following 4h of induction. Scale bars represent 1μm.

Spotting of the Δ*trpBA* mutant harbouring pSF251 on BDM, as well as growth analysis in liquid BDM, demonstrated full restoration of growth in the absence of tryptophan supplementation (Fig. 3B, C), confirming functional complementation of the auxotrophic phenotype. Next, we tested arabinose inducible expression using pSF251 by expressing the *S. acidocaldarius* S-layer protein SlaA-HA (21). Consistent with previous reports, basal expression of our protein of interest was present in the absence of induction, likely due to promoter leakiness (21, 22). Upon 4h induction with L-arabinose, an increase in SlaA-HA protein expression can be seen in strains grown in both BNS and BDM (Fig. 3D). Notably, lower expression levels were observed in BNS, potentially due to loss of the pSF251:SlaA-HA vector in the presence of tryptophan in the rich medium, which relieves selection pressure. Localization of SlaA-HA at the cell surface was also confirmed via immunofluorescence imaging (Fig. 3E). Together, these findings confirm that the tryptophan complementation system supports inducible protein expression and can be readily applied to cell biological studies in *S. acidocaldarius*.

### Dual-plasmid expression reveals co-enrichment of SlaA and SlaB at the division midzone

We previously reported that the addition of purified SlaA proteins to Δ*slaA S. acidocaldarius* cells arrested at division results in the accumulation of exogenous SlaA at the division bridge (21), likely due to the self-assembly of the lattice forming SlaA at this region. We wondered if this localization was mediated by the S-layer membrane anchor SlaB. As SlaB is an integral membrane protein, this could not be tested through the addition of purified protein to division arrested cells. Therefore, we co-transformed two plasmids into the Δ*trpBA* mutant, enabling the simultaneous inducible expression of the dominant negative Walker B mutant of the AAA-ATPase Vps4^E209Q^-His_6_ to enable the arrest of dividing cells, together with the S-layer proteins, utilizing both the *pyrEF* and *trpBAD* selection systems respectively (Fig. S4A).

Growth of double-positive cells in BDM was characterized by an extended lag phase and reduced expression levels, resulting in low signal intensity and poor representation of double-positive cells (results not shown). To improve growth and expression while maintaining selection stringency, cultures were subsequently propagated in Brock medium supplemented with casamino acids (BCS) instead of the dropout amino acid mix. Stringent selection of the Δ*trpBA* mutant was maintained in BCS medium (Fig. S4B), as casamino acids provides a rich but effectively tryptophan-free nutrient source due to its means of production via the acid hydrolysis of casein. Under these conditions, cells exhibited improved growth and substantially higher expression levels while maintaining stable co-expression of both constructs.

Immunoblot analysis confirmed the expression of all proteins after 8h of induction with L-arabinose in BCS (Fig. 4A). This was further supported by flow cytometric analysis (Fig. S4C), which also revealed the accumulation of a large population of cells containing greater than 2N DNA content following induction of Vps4^E209Q^-His_6_ expression (24), indicative of cell division arrest (Fig. S4C). While expression of the integral membrane protein SlaB was markedly lower than that of SlaA (Fig. 4A, S4C), this observation is consistent with previous challenges associated with heterologous expression of membrane proteins in *Sulfolobus*.

**Figure 4.**
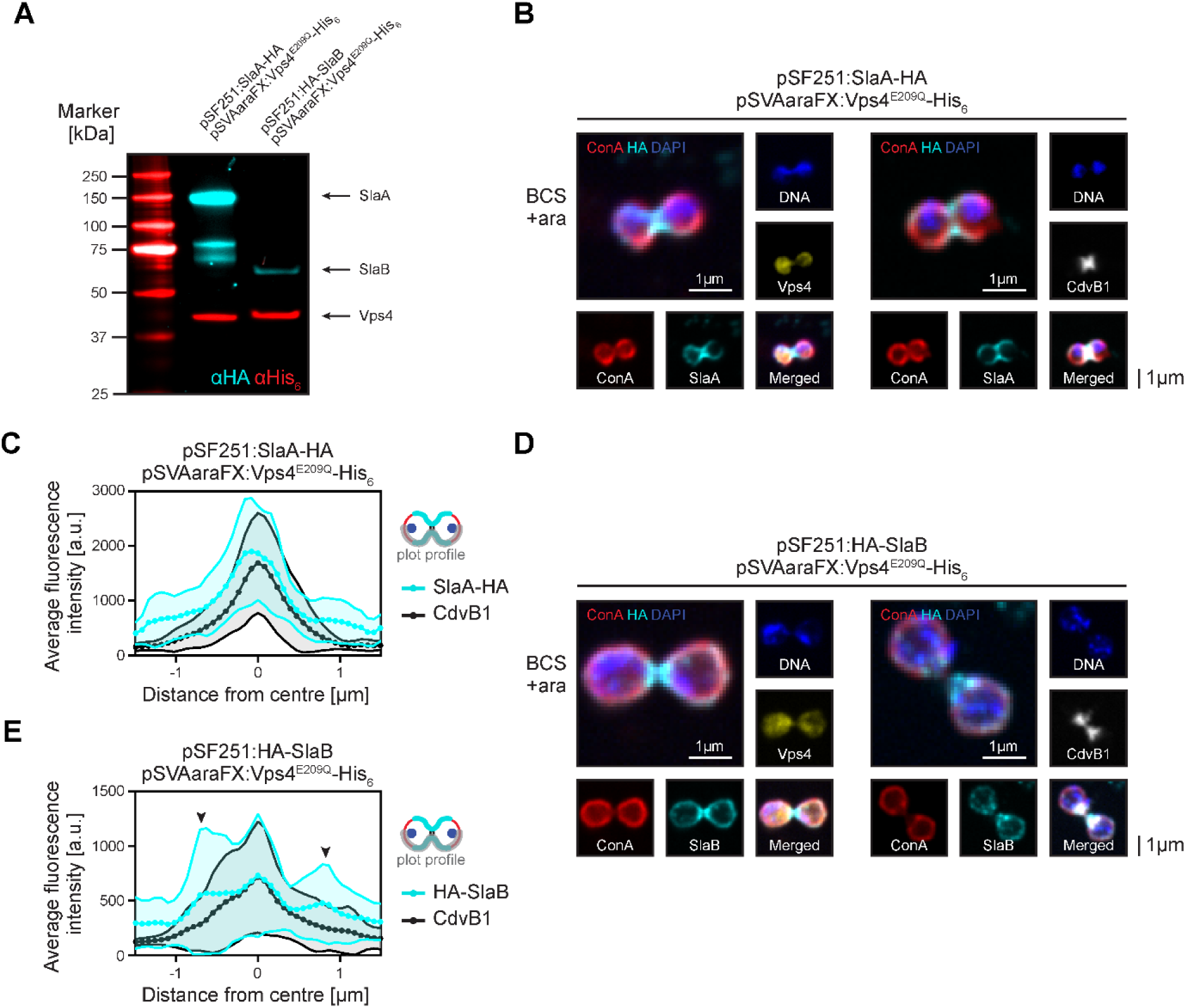
Dual expression of Vps4^E209Q^ and the S-layer proteins reveals accumulation of endogenous S-layer at the midzone of division arrested cells. **(A)** Immunoblot of a Δ*trpBA* strain expressing SlaA-HA or HA-SlaB (*trpBAD* selection) and Vps4^E209Q^-His_6_ (*pyrEF* selection) following 8h of arabinose induction in BCS. **(B)** Representative single-plane confocal micrographs of Δ*trpBA* in BCS expressing Vps4^E209Q^-His_6_ with SlaA-HA following 8h of arabinose induction. Cells were co-stained for HA with His_6_ (left) or CdvB1 (right). **(C)** Plot profile of SlaA-HA (cyan) and CdvB1 (black) fluorescence intensity across the division bridge marked by the cell division protein CdvB1; N=10 cells, shaded area represents mean ±SD. **(D)** Representative single-plane confocal micrographs of Δ*trpBA* in BCS expressing Vps4^E209Q^-His_6_ with HA-SlaB following 8h of arabinose induction. Cells were co-stained for HA with His_6_ (left) or CdvB1 (right). **(E)** Plot profile of HA-SlaB (cyan) and CdvB1 (black) fluorescence intensity across the division bridge marked by the cell division protein CdvB1; N=10 cells, shaded area represents mean ±SD. Black arrows represent the peaks of SlaB enrichment adjacent to the division bridges. All scale bars represent 1μm.

Confocal immunofluorescence imaging revealed enrichment of SlaA-HA at the midzone of cells arrested in division by Vps4^E209Q^-His_6_ (Fig. 4B, C), consistent with previous reports utilizing exogenously supplied purified SlaA (21). HA-SlaB was distributed around the cell periphery but also showed modest enrichment at the division bridge in a subset of arrested cells (Fig. 4D, E). Quantification of fluorescence intensity profiles confirmed the preferential accumulation of both SlaA and SlaB at the division midzone marked by the cell division protein CdvB1, although enrichment of SlaB was considerably weaker than that observed for SlaA (Fig. 4C, E). Interestingly, cells possessing long and narrow division bridges occasionally exhibited HA-SlaB enrichment proximal to the cell bodies, with reduced signal within the bridge itself (Fig. 4D, right), helping to explain the localisation pattern of exogenous SlaA seen previously (21). Consequently, fluorescence intensity profiles of these cells displayed two maxima flanking the division midzone (Fig. 4E, black arrows). This reveals that SlaA and SlaB have different localisation patterns, and raises the possibility that the spatial distribution of SlaB changes during bridge maturation.

Taken together, the localization of HA-SlaB at the division bridge suggests that components of the S-layer are enriched at this site during cytokinesis. Moreover, the co-enrichment of SlaA and SlaB at the division midzone is consistent with the possibility that membrane-associated SlaB contributes to the accumulation or stabilization of SlaA at this region, providing a potential explanation for the previously observed localization of exogenously supplied SlaA in division-arrested cells (21). This local deposition of S-layer material at the bridge, is then able to function in division to accelerate cytokinesis (21). These observations were made possible through the dual-marker system developed here, illustrating its utility for probing complex cell biological questions in *S. acidocaldarius*.

## Discussion

In this study, we expand the genetic toolkit available for *S. acidocaldarius* by establishing tryptophan auxotrophy as a robust and versatile selectable marker. Central to this development is the formulation of a defined Brock-based dropout medium BDM, which enables stringent selection by eliminating background growth associated with complex substrates. This overcomes a key limitation that has historically hindered the use of amino acid auxotrophy in this organism. This approach also opens the possibility of expanding the repertoire of selectable markers based on amino acid auxotrophy, such as leucine and histidine, which are commonly used selectable markers in model organisms including *S. pombe* (23, 25).

Using two markers, it becomes possible to co-transform and maintain two independently selectable plasmids to study the simultaneous expression of multiple proteins of interest, facilitating experiments combining fluorescent reporters with genetic or biochemical perturbations. This is useful because while chromosomal knock-ins are possible in *S. acidocaldarius* (12), they are often laborious, locus-dependent, and less flexible for rapid experimental iteration. Furthermore, plasmid-based expression systems allow faster construction and straightforward mixing and matching of promoters, tags, and reporter constructs, greatly increasing experimental flexibility.

Although growth in defined BDM was associated with a longer lag phase and lower arabinose-inducible protein expression than in rich BNS medium, BDM is primarily intended as a selective medium rather than a replacement for routine cultivation. Strains and plasmids can be readily selected under defined conditions and subsequently propagated in BNS for downstream experiments, such as *pyrEF*-based overexpression experiments requiring higher protein yields. Such approaches may prove valuable for applications including conditional expression systems and transposon-based genetic screens coupled with plasmid rescue strategies. In contrast, Brock medium supplemented with casamino acids provides a more complex nutrient environment while remaining effectively devoid of tryptophan, thereby maintaining stringent selection of the *trpBA* system. These observations suggest that future studies employing additional auxotrophic markers, such as histidine or leucine biosynthesis genes, may similarly require optimization of growth conditions to balance selection stringency with physiological robustness and expression levels. It will also be of interest to determine whether alternative chemically defined formulations (11) can provide improved growth characteristics while retaining the advantages of precisely controlled culture conditions.

We were unable to utilize the toxic antimetabolite 5-FAA as a counterselection marker in *S. acidocaldarius*. Deletion of the anthranilate isomerase *trpF*, the homolog of *TRP1* in *Saccharomyces cerevisiae* (23), was insufficient to render the mutant resistant to 5-FAA toxicity. This observation is consistent with previous findings in *Pyrococcus furiosus* (26). One possible explanation is that the elevated growth temperature of these thermophilic archaea may influence 5-FAA stability, uptake, or metabolism, resulting in toxicity independent of TrpF activity. Alternatively, an unknown mechanism or novel biosynthetic pathway underlying 5-FAA toxicity in these archaea may account for this phenotype.

Finally, we demonstrated the utility of dual auxotrophy selection in *S. acidocaldarius* through the co-transformation and simultaneous expression of two independently selectable plasmids in BCS medium. By combining expression of the dominant-negative Walker B mutant Vps4^E209Q^, which arrests cytokinesis, with epitope-tagged S-layer proteins, we were able to visualize the accumulation of SlaA and SlaB at the division bridge. The co-enrichment of SlaA and SlaB at the division midzone is consistent with a model in which SlaB contributes to the recruitment or stabilization of SlaA during cytokinesis as seen previously through the exogenous addition of SlaA (21), although further experiments will be required to establish the mechanistic relationship between these proteins. These observations suggest that local S-layer assembly may be spatially coordinated with division to aid cytokinesis in *S. acidocaldarius* (21), and demonstrate how dual auxotrophic selection can rapidly facilitate exploratory cell biological investigations prior to more laborious chromosomal engineering approaches.

Together, the development of a defined amino acid auxotrophy selection system and dual-selection genetic manipulation substantially broadens the experimental toolkit available for *S. acidocaldarius*. These advances provide increased flexibility in strain construction and cell biological studies combining genetic perturbations with reporter constructs. More broadly, they establish a platform for investigating how cellular structures are assembled and coordinated during archaeal growth and division, further strengthening the utility of this archaeon as a model system for archaeal cell biology and evolution.

## Materials and Methods

### Strains and growth conditions

*S. acidocaldarius* strains were grown in Brock medium (1) supplemented with 0.2% (w/v) sucrose, and further supplemented with either 0.1% (w/v) NZ-amine (BNS), an amino acid mix (Fig. 2A) (BDM), or 0.2% (w/v) casamino acids (BCS). All liquid medium were adjusted to pH 3.0 using sulfuric acid. All strains grown in liquid medium were kept at 75°C with constant shaking. Cultures were supplemented with 4mg/L uracil and/or 50mg/L tryptophan as required, unless otherwise stated. Solid medium was prepared by mixing in a 1:1 ratio, pre-heated 1.2% gelrite (w/v) with twice concentrated pH 5.0 medium, supplemented with 20mM CaCl_2_. All strains used in this study are shown in Table 1.

**Table 1:** Strains used in this study.

| Strain | Genotype | Figures | Source |
| --- | --- | --- | --- |
| MW001 | $\Delta pyrE$ | 1A-C, 2B-C, S2A, 3B, S4B | (12) |
| SF325 | $\Delta pyrE \Delta trpBA$ | 1A-C, 2B-D, S2B, 3B-C, S4A-C | This study |
| SF57 | $\Delta pyrE pSVAaraFX:slaA-HA$ | S2C-D | (21) |
| SF360 | $\Delta pyrE \Delta trpBA pSF251$ | 3B-C | This study |
| SF393 | $\Delta pyrE \Delta trpBA pSF251:slaA-HA$ | 3D-E | This study |
| SF469 | $\Delta pyrE \Delta trpF$ | S3A-C | This study |
| SF418 | $\Delta pyrE \Delta trpBA$<br>$pSF251:slaA-HA$<br>$pSVAaraFX:vps4^{E209Q-His_6}$ | 4A-C, S4A, S4C | This study |
| SF441 | $\Delta pyrE \Delta trpBA$<br>$pSF251:HA-slaB$<br>$pSVAaraFX:vps4^{E209Q-His_6}$ | 4A, 4D-E, S4A, S4C | This study |

Frozen stocks were made by resuspending cells in BNS or BDM containing 50% glycerol (v/v). *S. acidocaldarius* stocks were thawed by scraping with a sterile pipette tip and inoculating in 15mL of room temperature liquid medium. All thawed strains were grown for one to three days, and then passaged overnight to a final OD_595_ of 0.10 to

0.25 at the start of all experiments. Protein expression was induced by the addition of L-arabinose a final concentration of 0.2% (w/v) in exponentially growing cultures.

5-fluoroanthranilic acid and 5-fluoroorotic acid were prepared at a stock concentration of 100mg/mL in DMSO. Amino acid mix was prepared as a 20x stock solution in water (Fig. 2A), autoclaved, aliquoted and stored at −20°C. Wolin’s vitamin solution (20) was prepared as a 100x stock solution with 2mg/L biotin, 2mg/L folic acid, 10mg/L pyridoxine hydrochloride, 5mg/L thiamine-HCl, 10mg/L riboflavin, 5mg/L nicotinic acid, 5mg/L pantothenic acid, 0.1mg/L vitamin B12, 5mg/L aminobenzoic acid, and 5mg/L lipoic acid in water. The solution was autoclaved and stored at 4°C. Uracil was prepared as a 0.4g/L stock solution in water. Tryptophan stock solution was prepared at a concentration of 5g/L in water and kept in the dark to reduce oxidative degradation. Fresh stock was prepared upon observable oxidation or browning of the solution. All stock solutions were prepared in ultrapure water and filtered sterilized unless otherwise stated.

### Molecular genetics

The Δ*trpBA* tryptophan auxotrophic strain SF325 was constructed via the classical pop-in/pop-out method with plasmid pSVA406 (12). A double-stranded DNA fragment comprising 612bp upstream of *trpB* (*saci1422*) start codon and 612bp downstream of the *trpA* (*saci1421*) stop codon, with ApaI and NdeI restriction sites appended to the 5’ and 3’ ends respectively, was custom synthesized (IDT gBlock). This was cloned into pSVA406 using standard restriction digestion and ligation methods (5). For the Δ*trpF* strain, the *trpF* (*saci1424*) open reading frame was deleted from the stop codon of the *trpD* gene (*saci1423*) to the start codon of the *trpE* gene (*saci1425*) to avoid disruption of the overlapping up and downstream genes. Homology regions of 487bp up and downstream of the targeted region were used as described above.

HA-SlaB construct was designed as described previously (21), with the HA tag (YPYDVPDYA) inserted downstream of the signal sequence (amino acid residue S24) of SlaB. The tryptophan complementation vector pSF251 (Fig. 3A) was constructed by replacing the *pyrEF* cassette of the arabinose-inducible expression plasmid pSVAaraFX-HA (22) with the *trpBAD* genes of *Sc. solfataricus* (*sso0888* - *sso0890*) between the KpnI and XmaI restriction sites. All plasmids used in this study are summarised in Table 2.

**Table 2:**
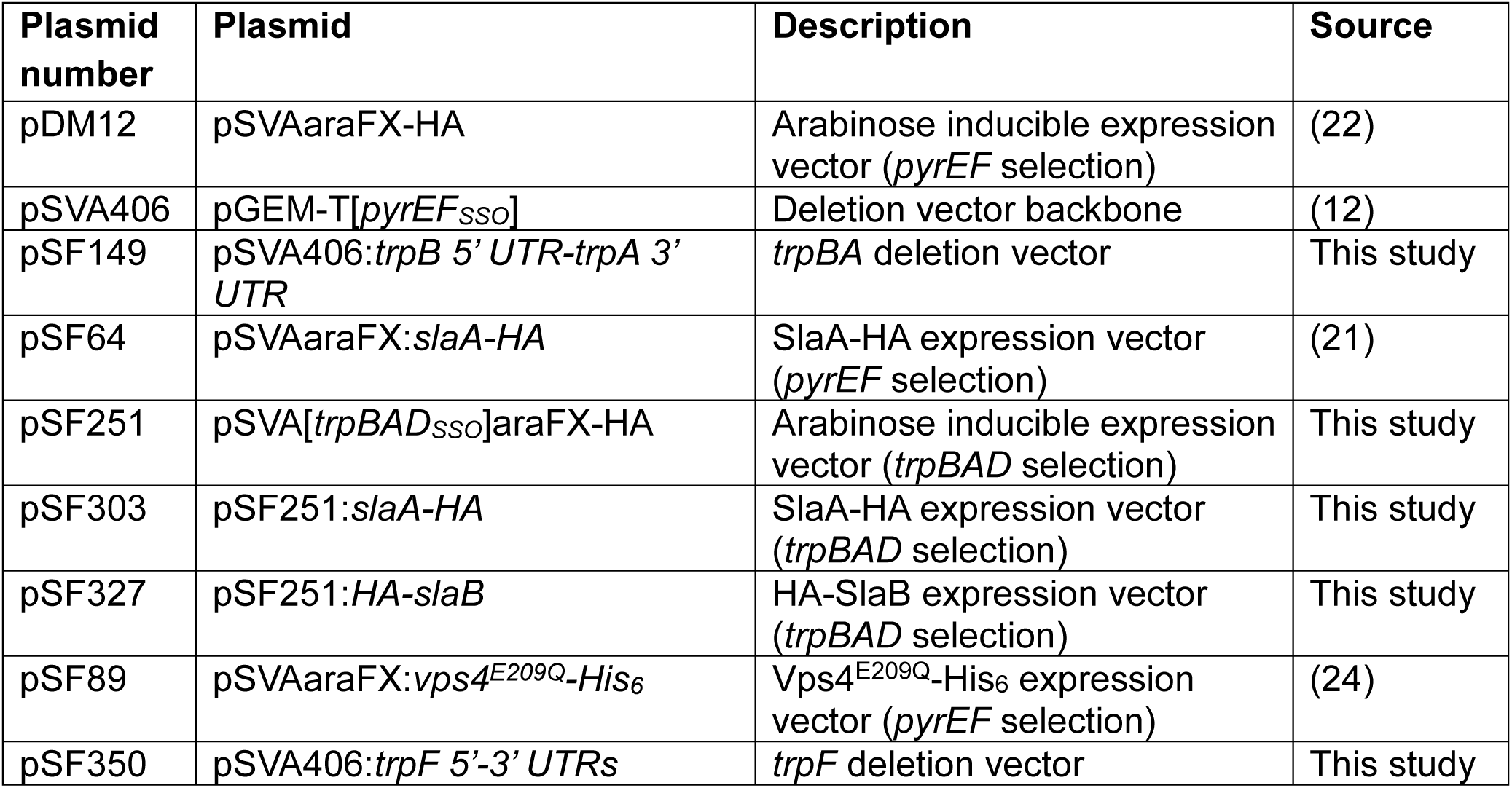
Plasmids used in this study.

| Plasmid number | Plasmid | Description | Source |
| --- | --- | --- | --- |
| pDM12 | pSVAaraFX-HA | Arabinose inducible expression vector (*pyrEF* selection) | (22) |
| pSVA406 | pGEM-T[*pyrEF*<sub>SSO</sub>] | Deletion vector backbone | (12) |
| pSF149 | pSVA406:*trpB* 5' UTR-*trpA* 3' UTR | *trpBA* deletion vector | This study |
| pSF64 | pSVAaraFX:*slaA*-HA | *SlaA*-HA expression vector (*pyrEF* selection) | (21) |
| pSF251 | pSVA[*trpBAD*<sub>SSO</sub>]araFX-HA | Arabinose inducible expression vector (*trpBAD* selection) | This study |
| pSF303 | pSF251:*slaA*-HA | *SlaA*-HA expression vector (*trpBAD* selection) | This study |
| pSF327 | pSF251:HA-*slaB* | HA-*SlaB* expression vector (*trpBAD* selection) | This study |
| pSF89 | pSVAaraFX:*vps4*<sup>E209Q</sup>-His<sub>6</sub> | *Vps4*<sup>E209Q</sup>-His<sub>6</sub> expression vector (*pyrEF* selection) | (24) |
| pSF350 | pSVA406:*trpF* 5'-3' UTRs | *trpF* deletion vector | This study |

Quick genomic DNA extraction from *S. acidocaldarius* and *Sc. solfataricus* for PCR amplification of genes of interest or PCR genotyping was performed as follows. 500µL of exponentially growing cells were collected via centrifugation at maximum speed for 30s. The cell pellet was resuspended in 20µL of 0.2M NaOH to lyse them, and then diluted with 100µL of 0.2M Tris-HCl pH 8.0.

Preparation of electrocompetent *Sulfolobus* cells and transformation via electroporation were performed as described previously (5). For strains containing two plasmids with different auxotrophic selection, transformation was performed using equal amounts of both methylated plasmids (approximately 200ng each) in a single electroporation reaction, and selected on BDM-gelrite plates without uracil or tryptophan supplemented. After 7-10 days of incubation at 75°, isolated colonies were restreaked onto fresh BDM-gelrite plates and grown for a further two to three days, followed by colony PCR genotyping. Positive clones were then inoculated into liquid medium, grown for three days, and frozen as described above.

### Spot assays

*S. acidocaldarius* strains were grown in the appropriate media to an OD_595_ of 0.20-

0.30. 2 OD_595_ of exponentially growing cells were then collected via centrifugation at 8000rcf for 3min, resuspended in 1mL of liquid medium followed by a 10-fold dilution to achieve a final theoretical OD_595_ of 0.2. Five further 10-fold serial dilutions were then performed to obtain final OD_595_ values of 0.02, 0.002, 0.0002, 0.00002, and 0.000002. A total of 5µL of each dilution was spotted onto BNS, BDM, or BCS plates containing 0.6% gelrite, supplemented with uracil and/or tryptophan as required. Plates were incubated for 5 to 7 days at 75°C.

### Growth curve measurements

*S. acidocaldarius* were thawed and grown in the appropriate medium as described above. 2 OD_595_ of exponentially growing cells were collected via centrifugation at 8000rcf for 3min and resuspended in 1mL of fresh, pre-warmed media. 50µL of cells were then inoculated into 50mL of pre-warmed media, giving a theoretical starting OD_595_ of 0.002. OD_595_ readings were taken periodically over 5 to 7 days until stationary phase of cell growth.

### Immunostaining, confocal microscopy and flow cytometry

Cells were fixed by three stepwise additions of 4°C ethanol to a final concentration of 70%. 3mL of exponentially growing cells was added to 1.5mL of 4°C ethanol for 5min. This is followed by a further addition of 1.5mL of 4°C ethanol for 5min, before adding 4mL of 4°C ethanol to achieve a final concentration of 70% ethanol. Fixed cells were stored at 4°C for up to 3 months. Confocal microscopy and flow cytometry were performed using fixed cells as previously described (5, 21).

### Western blotting

1 OD_595_ of cells were pelleted and resuspended in 200 µL of 1x NuPAGE LDS sample buffer (Invitrogen, NP0007) containing 10% β-mercaptoethanol. The samples were boiled for 5min before loading onto a NuPAGE 4 to 12% Bis-Tris gel (Invitrogen, NP0321BOX) with a protein standard (BioRad, 1610373). SDS-PAGE was performed at a constant 150V for 65min in MES-SDS running buffer. Wet transfer of separated proteins to a nitrocellulose membrane was performed at a constant 100V for 1h at 4°C, following standard Western blotting protocols. Blocking and antibody incubation was performed in phosphate-buffered saline containing 0.2% Tween-20 (Sigma-Aldrich, P9416) and 5% milk, as described previously (5). Primary and secondary antibodies used are listed in Table 3.

**Table 3:**
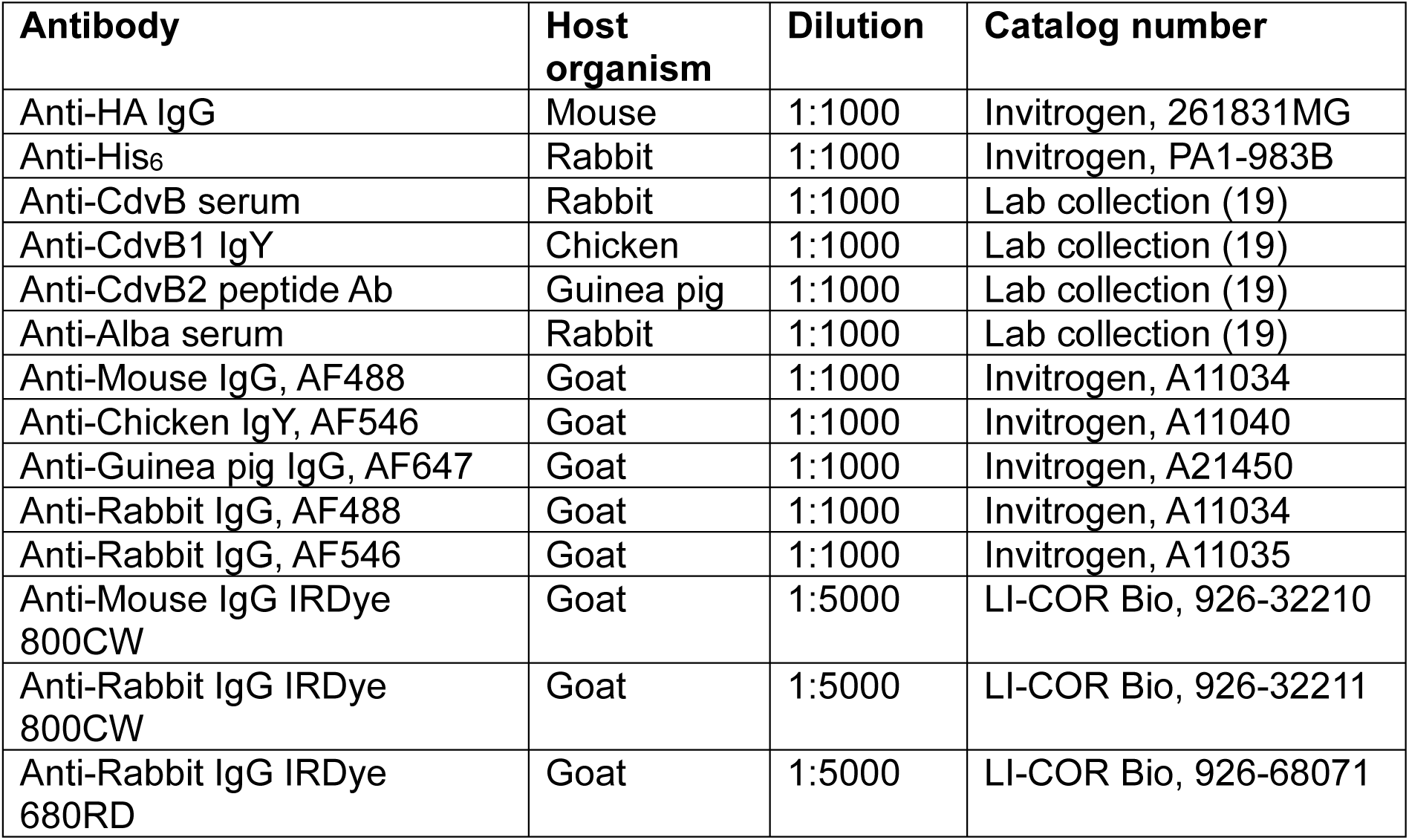
Antibodies used in this study.

### Quantification and statistical analyses

All statistical analysis was performed on Microsoft Excel or GraphPad Prism 10 software. All data presented are shown as mean ±SD, unless otherwise specified.

## Author contributions

S.F. conceived the project, performed all experiments and wrote the manuscript. B.B. supervised the project and co-edited the manuscript.

## Acknowledgements

We would like to thank Dr. Yin-Wei Kuo and Dr. Arthur Radoux-Mergault for helpful discussions on experimental design. We also thank Martyn Howard and the team at the Medical Research Council Laboratory of Molecular Biology Media Prep and Glasswash facility for their technical support in this work. All flow cytometry experiments were performed at the MRC-LMB Flow Cytometry facility.

S.F. was supported by the Wellcome Trust (222460/Z/21/Z); B.B. received support for work in *Sulfolobus* from the Medical Research Council- Laboratory of Molecular Biology (MC_UP_1201/27), the Wellcome Trust (222460/Z/21/Z). This work was supported by the Moore-Simons Project on the Origin of the Eukaryotic Cell, Simons Foundation (735929LPI).

## Data and materials availability

All data are available in the manuscript or the supplementary materials. All materials are available from the corresponding authors with a completed materials transfer agreement in compliance with institutional policy.

---

**Figure S1.**
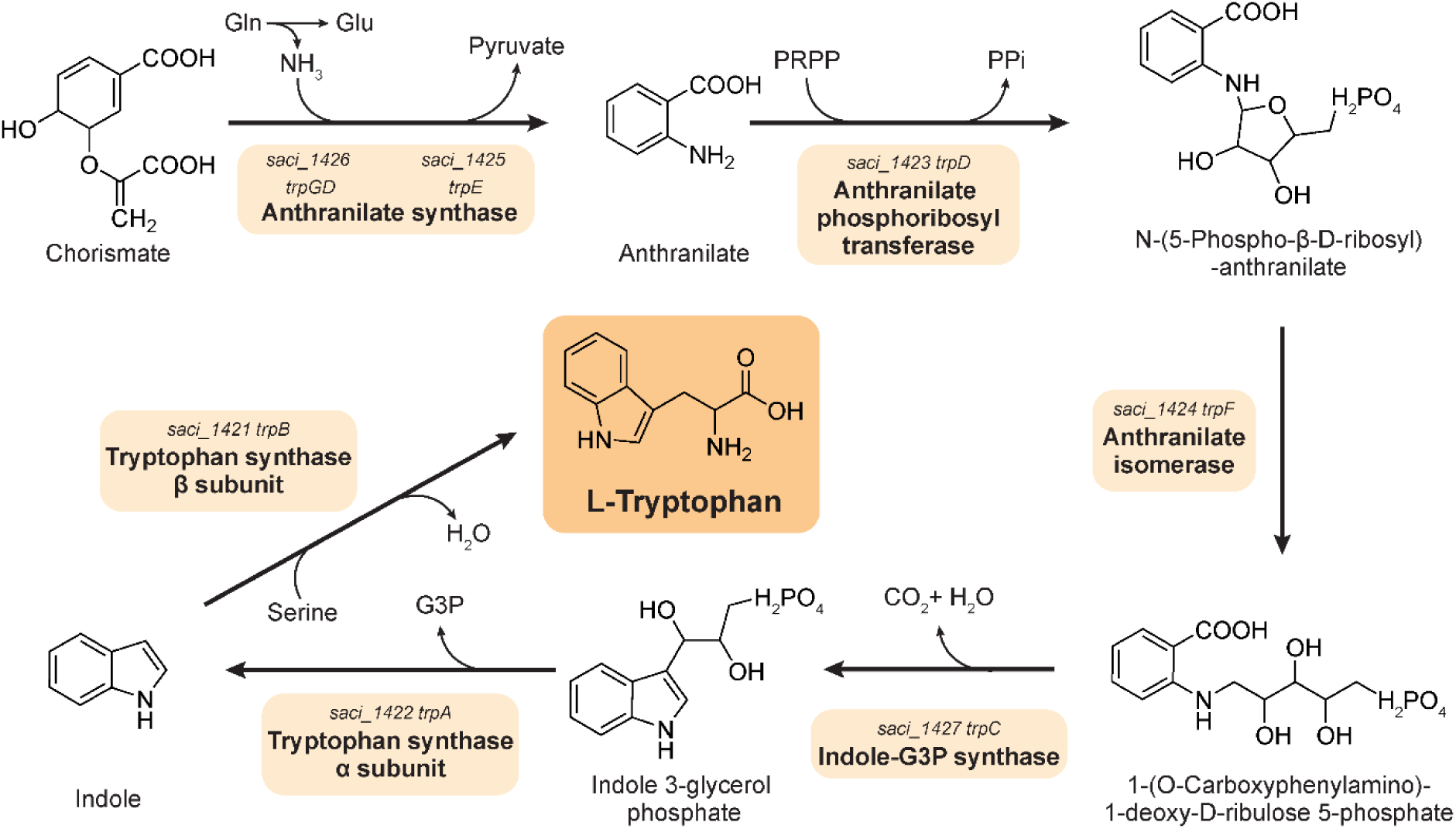
The tryptophan biosynthetic pathway. Tryptophan is synthesized from chorismate, a crucial intermediate in the synthesis of aromatic amino acids, vitamins and other metabolites. It is first converted to anthranilate via anthranilate synthase (TrpE/TrpGD). Anthranilate is then converted to N-(5-phosphoribosyl)-anthranilate by the anthranilate phosphoribosyl transferase (TrpD), followed by isomerization to 1-(o-carboxyphenylamino)-1-deoxyribulose-5-phosphate by phosphoribosylanthranilate isomerase (TrpF). This intermediate is subsequently cyclized to indole-3-glycerol phosphate by indole-3-glycerol phosphate synthase (TrpC). In the final steps, tryptophan synthase, a heterotetrameric complex composed of TrpA and TrpB subunits, catalyzes the conversion of indole-G3P into indole, before its condensation with serine into tryptophan.

**Figure S2.**
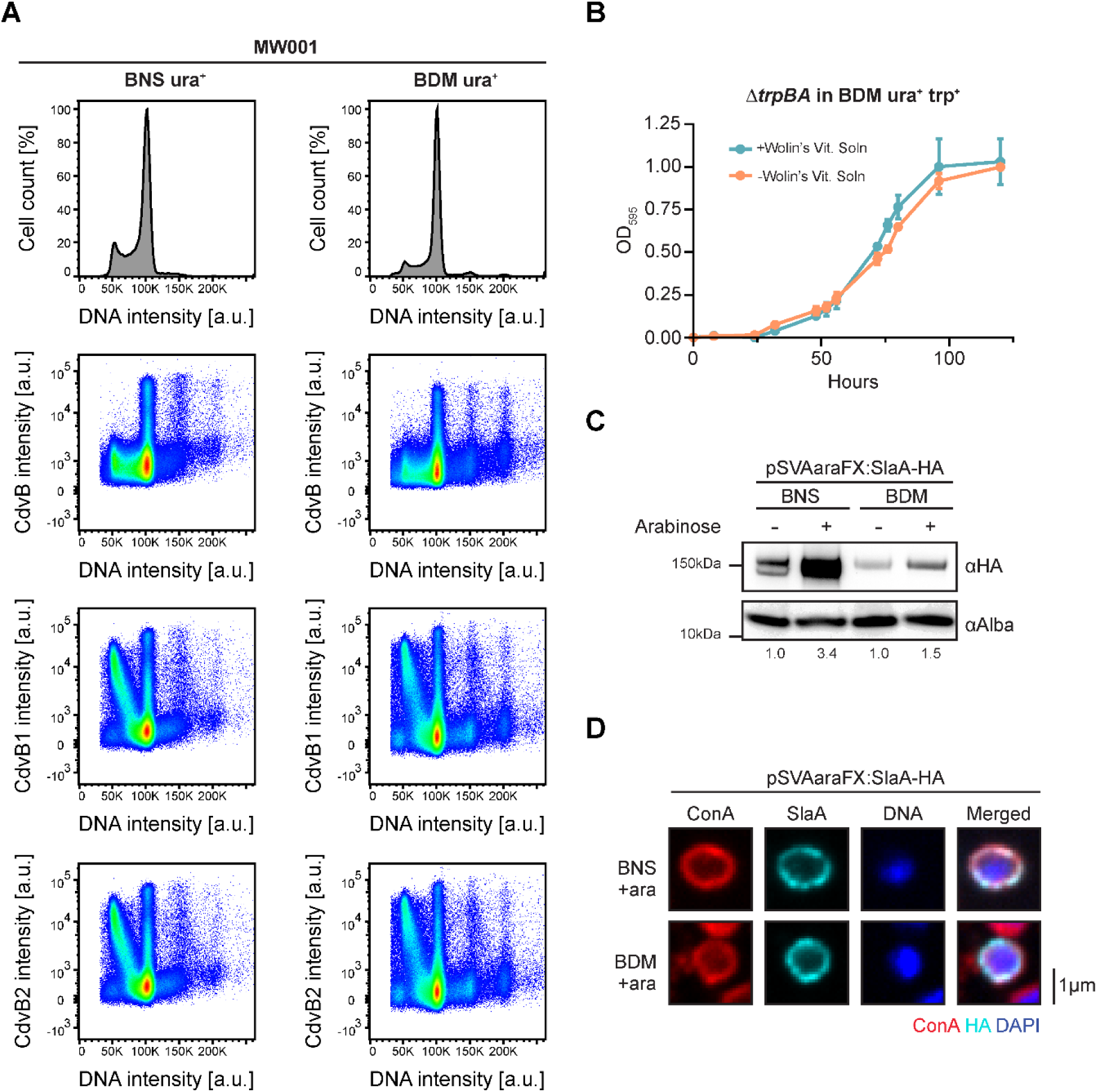
Characterization of *S. acidocaldarius* growth in BDM. **(A)** Representative flow cytometry analysis of MW001 grown in BNS (left) or BDM (right) showing DNA content and cell cycle-dependent profiles of ESCRT-III homologs CdvB, CdvB1, and CdvB2. Comparable distribution patterns were observed between the two growth conditions. Each spot represents a single cell, with the density gradient going from blue to red (N=3, n=0.5×10^6^ events). **(B)** Growth analysis of MW001 in BDM with or without supplementation of Wolin’s vitamin solution (20), demonstrating a modest increase in growth rates in the linear phase upon vitamin addition. Error bars represents mean ±SD. **(C)** Immunoblot analysis of SlaA-HA expression following induction with arabinose for 4 h in cells grown in BNS and BDM using the pSVAaraFX-HA plasmid (*pyrEF* selection). The DNA-binding protein Alba was used as a loading control. Values indicate the normalised HA signal intensity expressed relative to the corresponding uninduced sample. **(D)** Representative single-plane confocal micrographs of cells expressing SlaA-HA, grown in BNS or BDM, following 4h of arabinose induction. Scale bar represents 1μm.

**Figure S3.**
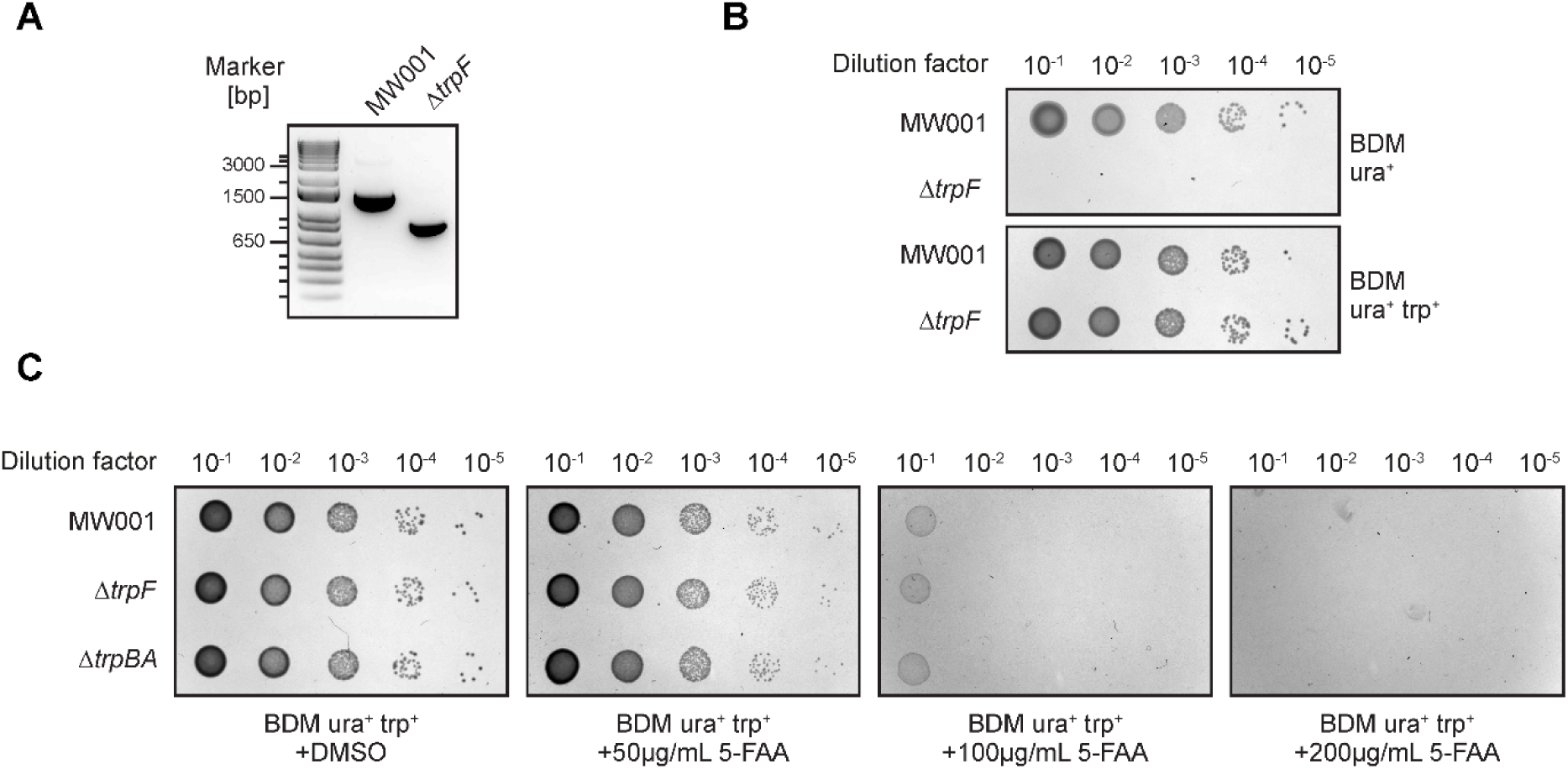
5-Fluoroanthranilic acid counterselection is ineffective in *S. acidocaldarius*. **(A)** PCR genotyping utilizing flanking primers confirms the deletion of the 600bp Δ*trpF* gene (*saci1424*) in the Δ*pyrE* MW001 background strain. **(B)** Spot assay showing tight selection of the Δ*trpF* mutant on BDM, with no background growth in the absence of tryptophan supplementation. **(C)** Spot assay of the indicated strains on BDM supplemented with uracil and tryptophan, in the presence of (left to right): DMSO control, 50μg/mL, 100μg/mL or 200μg/mL 5-fluoroanthranilic acid (5-FAA).

**Figure S4.**
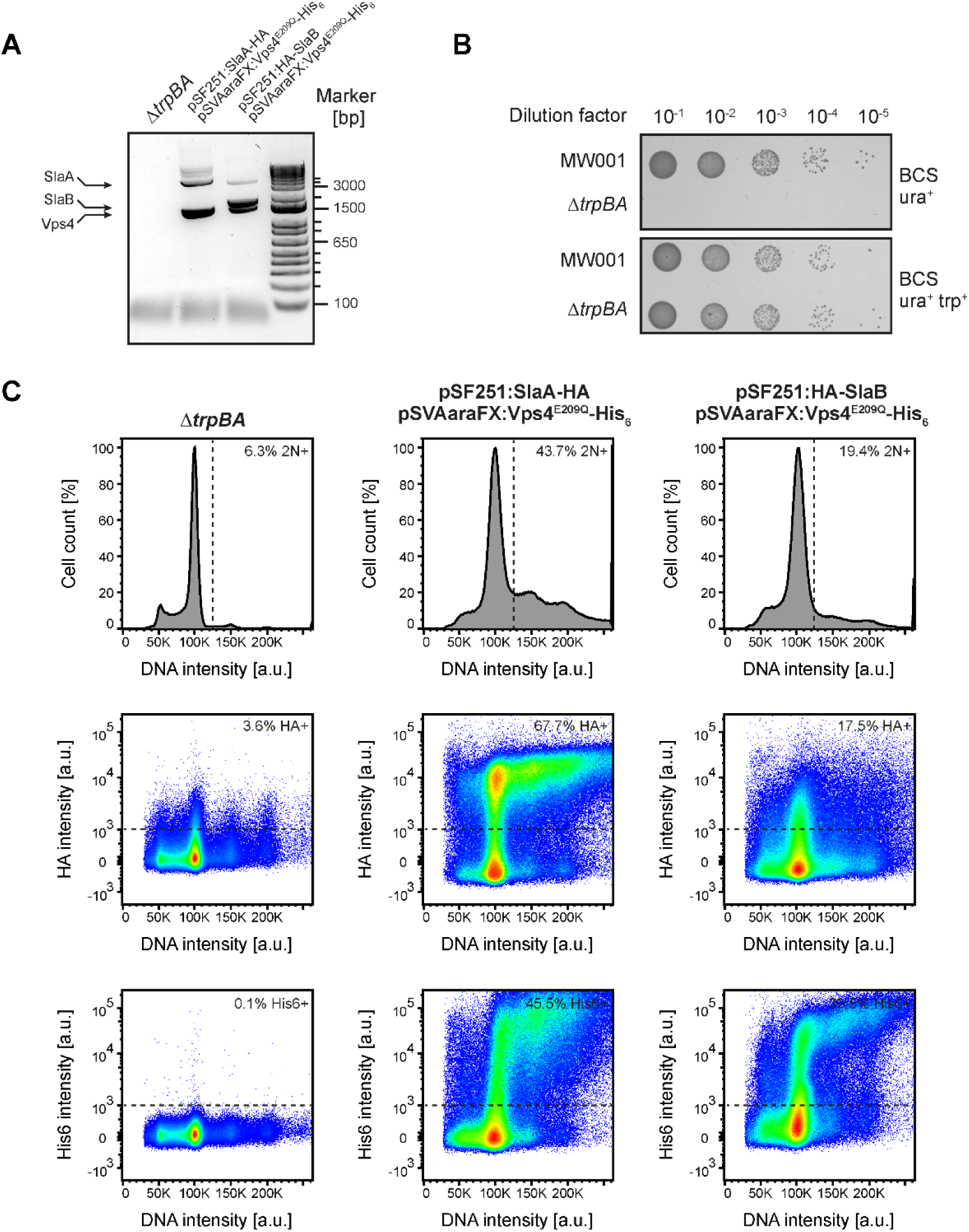
Co-transformation of Vps4^E209Q^ and the S-layer proteins. **(A)** PCR genotyping of the Δ*trpBA* strain and derivative strains co-transformed with plasmids expressing SlaA-HA and Vps4^E209Q^-His_6_ or HA-SlaB and Vps4^E209Q^-His_6_. Genotyping was performed using primers flanking the inserted expression cassettes. **(B)** Spot assay showing tight selection of the Δ*trpBA* mutant on Brock medium supplemented with casamino acids (BCS), with no background growth in the absence of tryptophan supplementation. **(C)** Representative flow cytometry analysis of Δ*trpBA* and derivative strains co-transformed with plasmids expressing SlaA-HA and Vps4^E209Q^-His_6_ or HA-SlaB and Vps4^E209Q^-His_6_ showing DNA content and scatter plots of HA and His_6_. Strains were grown in BCS and induction performed for 8h. Each spot represents a single cell, with the density gradient going from blue to red (N=3, n=0.5×10^6^ events). Dotted lines indicate the gating threshold used.

